# Spike-based phylogenetically defined clades within the *Alphacoronavirus 1* species

**DOI:** 10.1101/101774

**Authors:** Gary R. Whittaker, Nicole M. André, Jean Kaoru Millet

**Affiliations:** Department of Microbiology and Immunology, College of Veterinary Medicine, Cornell University, Ithaca NY 14853, United States

## Abstract

Taxonomic classification for the *Coronaviridae* can be challenging, due to the wide host tropism and highly variable genome of the viruses in this Family. Within the *Alphacoronavirus* genus, there is a single species *Alphacoronavirus 1* that encompasses several biologically distinct viruses of distinct animal species. Here, we carried out phylogenetic analysis of members of the *Alphacoronavirus* genus, focusing on the viral spike gene, which is a primary driver of viral tropism and pathogenesis. We identify two distinct clades (A and B) within the *Alphacoronavirus 1* species. *Alphacoronavirus 1* clade A encompasses serotype I FCoV and CCoV, and *Alphacoronavirus 1* clade B, encompasses serotype II FCoV and CCoV and TGEV-like viruses. We propose this clade designation, along with the newly proposed *Alphacoronavirus 2* species, as an improved way to classify the diverse *Alphacoronavirus* genus.

## Introduction

Members of the *Coronaviridae* family form a diverse group of enveloped, single-strand, positive-sense RNA viruses. The *Coronaviridae* family is divided into the *Torovirinae* and *Coronavirinae* subfamilies, both of which contain viral species characterized by their exceptionally large RNA genomes in the 26.4-31.7 kb and 26.6-28.5 kb ranges, respectively. *Torovirinae* and *Coronavirinae* subfamilies members are able to infect a diverse array of vertebrate species. They have a distinct genomic architecture and replication strategy shared with other members of the *Nidovirales* order, which also includes the *Arteriviridae*, *Mesoniviridae*, and *Roniviridae* families (1, 2). Coronaviruses (CoV) are classified into four genera, with *Alphacoronavirus* and *Betacoronavirus* containing members that infect mostly mammalian species, and *Gammacoronavirus* and *Deltacoronavirus* grouping viruses infecting both birds and mammals (3). The zoonotic emergence of highly pathogenic human betacoronaviruses, severe acute respiratory syndrome coronavirus (SARS-CoV) in 2002, and Middle East respiratory syndrome coronavirus (MERS-CoV) in 2012, have renewed interest for the study of coronaviruses and highlighted their propensity for recombination and to cross the species barrier (4, 5).

The *Alphacoronavirus* genus is composed of viruses infecting bats, ferrets, mink, cats, dogs, pigs, and humans. The prototypical species *Alphacoronavirus 1* is composed of the following prototypical virus strains (6): feline coronavirus (FCoV), canine coronavirus (CCoV), and transmissible gastrointestinal enteric virus (TGEV). FCoV is an *Alphacoronavirus* of particular interest as it manifests as two distinct biotypes or pathotypes with a highly transmissible form, feline enteric coronavirus (FECV) that provokes self-limiting, usually mild enteric tract infections, and a systemic form, feline infectious peritonitis virus (FIPV), typically associated with low transmissibility but high morbidity (7). In the widely accepted “internal mutation” hypothesis, it is believed that genetic mutations in the genome of FCoV occur within an infected animal, giving rise to FIPV (8). A similar FIP-like pathogenesis phenomenon is also observed with ferret coronaviruses (FRCoV) (9). While CCoV is a widespread enteric virus of dogs and can occur in highly pathogenic forms, the virus does not manifest itself with FIP-like clinical signs (10).

The coronavirus spike (S) envelope glycoprotein, the main determinant of virus entry, is an essential structural protein as it governs binding to the host cell receptor, mediates viral membrane fusion, and is typically proteolytically processed by host cell proteases to activate its fusogenicity (11, 12). As such, the coronavirus S protein is a crucially important viral component as it determines to a large extent host species, tissue, and cell tropism as well as pathogenicity and transmission. Previous serological characterizations of alphacoronaviruses, based on the antigenicity of the S glycoprotein, have revealed the existence of two FCoV serotypes (serotype I and II) (13–15). Both serotype I and II FCoV can manifest as either FECV and FIPV biotypes. FCoV serotype I is more prevalent in cats than serotype II, but has proved more difficult to culture *in vitro* (16, 17). Similarly in strains of CCoV, a common enteric virus of dogs, two serotypes (CCoV I and II) have also been characterized, and are distinguished by genetic differences in the S and ORF3 genes. Serotype II CCoV strains can be further subdivided into the IIa, IIb, and IIc subtypes (10). CCoV IIa and IIb strains are distinguished by differences in the N-terminal domain of the S protein (NTD), where the IIb NTD is closely related to TGEV NTD. The recently characterized IIc subtype of CCoV has been reported in Sweden and in the United States.

The evolution of strains within the *Alphacoronavirus 1* species is complex and likely involved a number of recombination events. It is thought that serotype I FCoV and CCoV originated from a common ancestor. A recombination event occurring between a serotype I CCoV with an unknown coronavirus gave rise to serotype II CCoV which acquired a recombinant S protein, distinct from serotype I S envelope glycoprotein. TGEV appears to have originated from a serotype II CCoV (18). Additional, independent recombination events between serotype I FCoV and serotype II CCoV gave rise to serotype AI FCoV, such as FIPV WSU-79-1146 and FECV WSU-79-1683, which acquired a serotype II CCoV S protein (19). Furthermore, we have previously shown that CCoV strain A76 also has a recombinant S protein, a product of recombination between serotype I and II CCoV sequences (20). CCoV-A76 S was shown to have a serotype I-like NTD, while the rest of the protein was serotype II-like. Analysis of coronavirus recombination events within the S protein sequence revealed its modular nature, allowing exchanges of functional domains between co-infecting viruses (5, 20).

Because of the numerous recombination events occurring within the S gene of *Alphacoronavirus* 1 species, current classification of *Alphacoronavirus 1* strains are often not well defined and fail to recapitulate the previously established serotype demarcation. However, in addition to serological differences, serotype I and II S proteins are fundamentally different in several biological aspects. While the receptor for serotype II FCoV, CCoV, and TGEV has been shown to be aminopeptidase N (APN, or CD13), the receptor for serotype I strains remains unknown. Furthermore, the S protein of serotype I strains contain an additional cleavage site, the S1/S2 site, which is not present in serotype II or in other *Alphacoronavirus* S proteins. This site has been shown to be important for cell culture adaptation and pathogenesis of FCoV (21, 22). Because of the critical role played by the S glycoprotein in virus entry, pathogenesis, and tropism, and since the S proteins of serotype I and II strains differ greatly, we propose a classification of *Alphacoronavirus 1* strains into two clades (*Alphacoronavirus 1* clade A and B), using a functionally-based S protein sequence classification that reflects the previously determined serologically-based demarcation.

## Results

As a starting point to gain a better understating of the phylogenetic relationships between alphacoronaviruses, we generated a phylogenetic tree of key representative species and strains based on complete genome nucleotide sequence alignment, a method often used to classify coronaviruses (Fig. 1A). As expected, the *Alphacoronavirus 1* FCoV, CCoV, and TGEV strains formed a well-defined monophyletic group. The analysis also reveals the clearly delineated branching of coronavirus strains that infect ferrets and mink, FrCoV-NL-2010, MinkCoV-WD1127, and MinkCoV-WD1133, which were recently proposed to form a separate species, *Alphacoronavirus 2* (23) (Fig. 1A). Two main subgroupings within the *Alphacoronavirus 1* species were observed with CCoV and TGEV partitioning into one subgroup and FCoV forming a second, separate subgroup. However, as shown by the CCoV/FCoV serotype highlighting, the complete genome-based phylogenetic tree fails to cluster CCoV and FCoV strains according to serotype demarcations, and instead groups strains according to their host species.

**Figure 1.**
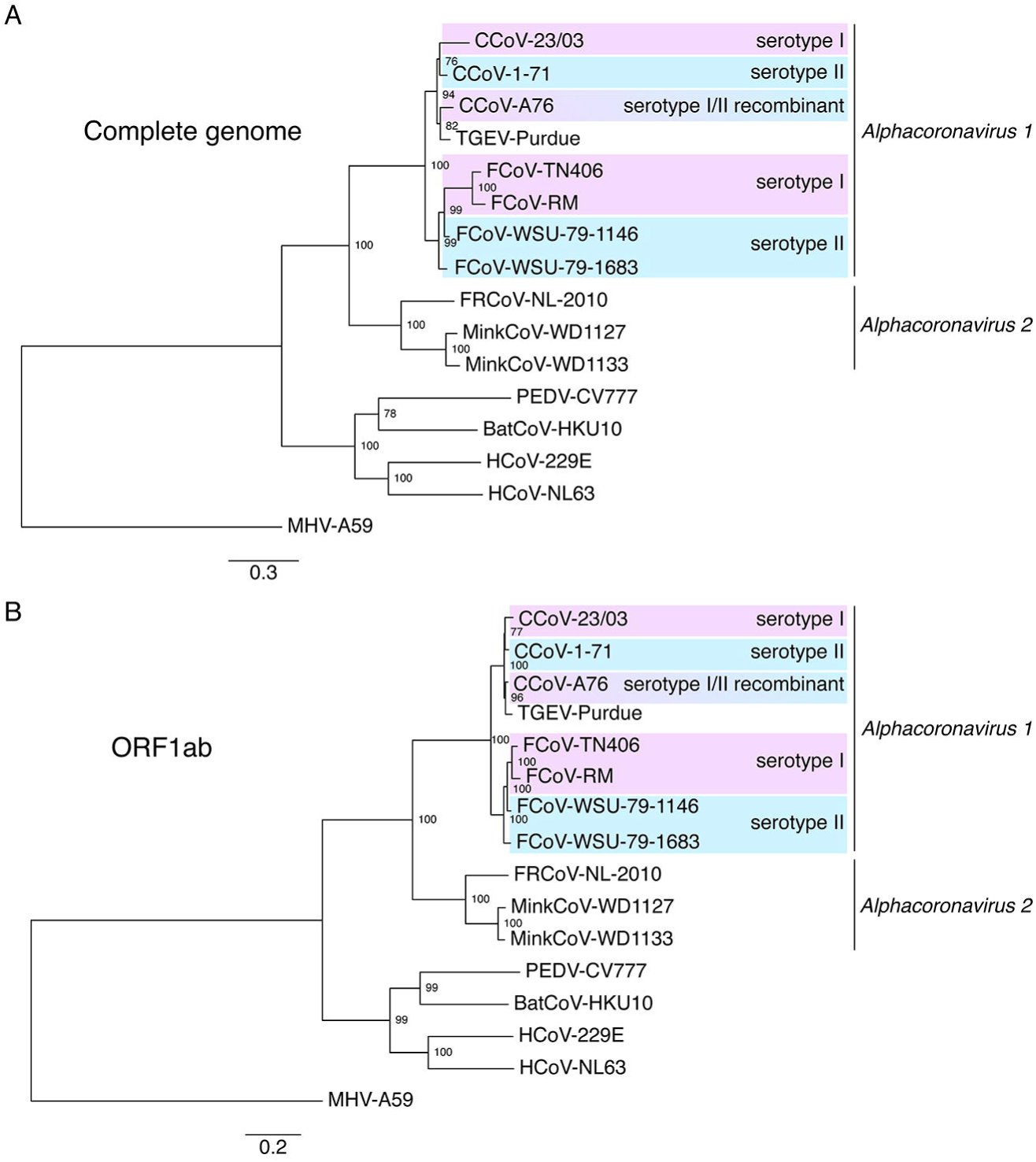
Phylogenetic analysis of alphacoronaviruses based on complete genome and ORF1ab protein sequence. Nucleotide sequences of the complete genomes of alphacoronaviruses and of the *Betacoronavirus* strain MHV-A59 (**A**) or the complete protein sequences of the ORF1ab polyprotein of the corresponding viruses (**B**) were aligned using MAFFT (http://mafft.cbrc.jp/alignment/software/) within Geneious 10 software package. The alignment was then used to generate a maximum-likelihood phylogenetic tree using PhyML (25). The tree was rooted with MHV-A59. Numbers at nodes indicate the bootstrap support on 100 replicates. Scale bar indicates the estimated number of substitutions per site. Accession numbers for complete genome nucleotide sequences used: CCoV-23/03 (KP849472.1), CCoV-1-71 (JQ404409.1), CCoV-A76 (JN856008.2), TGEV-Purdue (AJ271965.2), FCoV-TN406 (EU186072.1), FCoV-RM (FJ938051.1), FCoV-WSU-79-1146 (NC_002306.3), FCoV-WSU-79-1683 (JN634064.1), FRCoV-NL-2010 (NC_030292.1), MinkCoV-WD1127 (HM245925.1), MinkCoV-WD1133 (HM245926.1), PEDV-CV777 (AF353511.1), BatCoV-HKU10 (JQ989270.1), HCoV-229E (KU291448.1), HCoV-NL63 (AY567487.2), MHV-A59 (AY700211.1). Accession numbers for complete ORF1ab protein sequences used: CCoV-23/03 (AKZ66481.1), CCoV-1-71 (AFG19735.1), CCoV-A76 (AEQ61967.2), TGEV-Purdue (P0C6Y5.1), FCoV-TN406 (ABX60144.1), FCoV-RM (ACT10853.1), FCoV-WSU-79-1146 (YP_004070193.2), FCoV-WSU-79-1683 (AFH58022.1), FRCoV-NL-2010 (YP_009256195.1), MinkCoV-WD1127 (ADI80512.1), MinkCoV-WD1133 (ADI80522.1), PEDV-CV777 (P0C6Y4.1), BatCoV-HKU10 (AFU92103.1), HCoV-229E (AOG74782.1), HCoV-NL63 (AAS58176.2), MHV-A59 (AAU06353.1).

Analysis based on the ORF1ab polyprotein sequence, reveals very similar phylogenetic relationships within the *Alphacoronavirus 1* species, with a branching that partitions FCoV strains in one subgroup and CCoV and TGEV strains in another (Fig. 1B). In a similar manner than with the complete genome analysis, the ORF1ab polyprotein-based phylogenetic tree fails to group strains according to FCoV and CCoV serotypes.

One of the most distinguishing features found in serotype I FCoV and CCoV is a S protein cleavage site containing a polybasic furin recognition motif at the junction between the S1 receptor-binding domain and the S2 fusion domain (S1/S2, fig. 2A). While a furin site at S1/S2 is quite common in betacoronaviruses (e.g. MHV, HCoV-HKU1, and MERS-CoV) and gammacoronaviruses (e.g. IBV), it is not typically found in alphacoronaviruses. This observation highlighted the distinctive nature of serotype I S and prompted us to investigate more closely the phylogenetic relationships of *Alphacoronavirus 1* members by characterizing their S protein cleavage sites and performing S-protein-based phylogenetic analysis (fig. 2).

**Figure 2.**
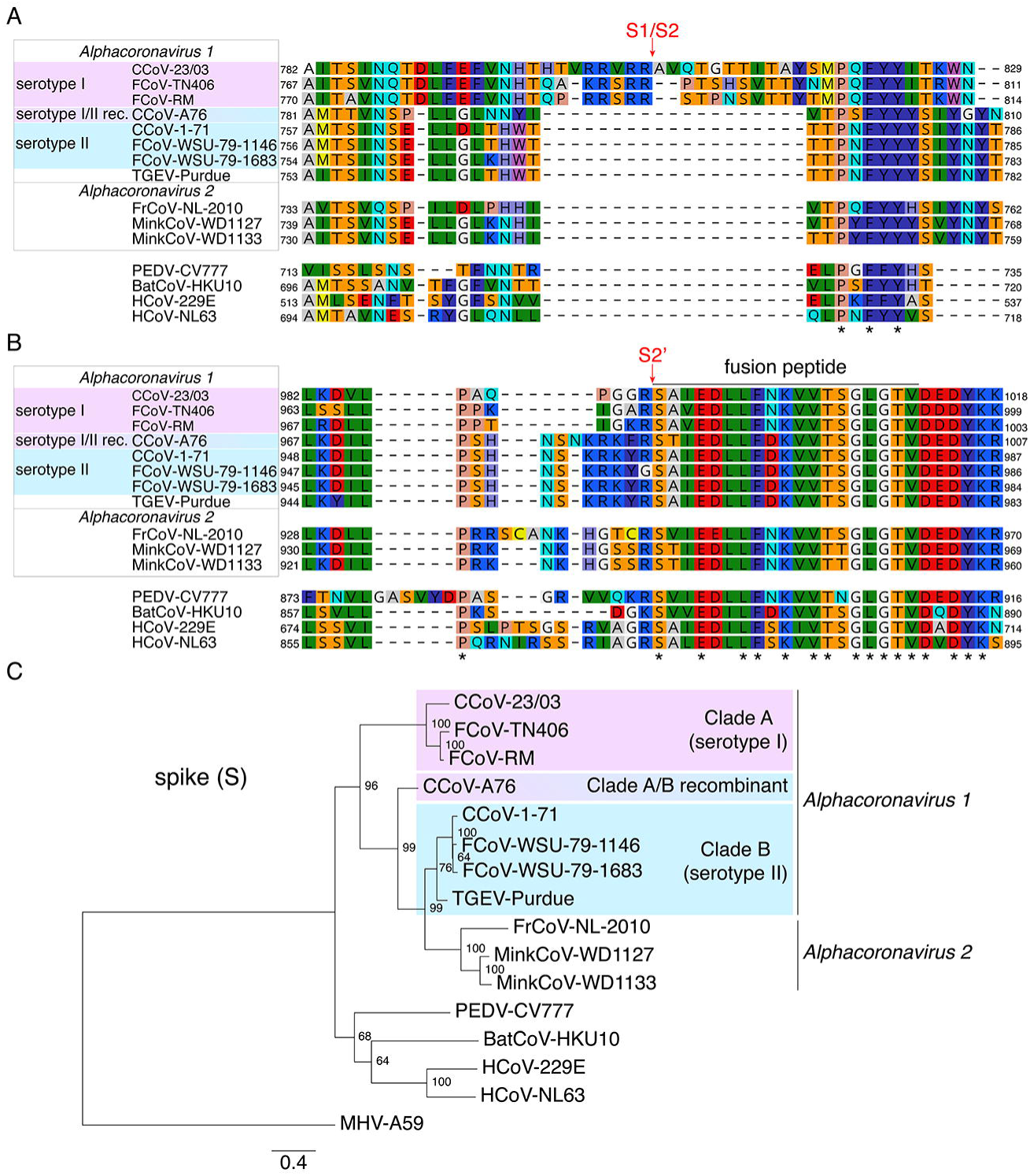
Analysis of *Alphacoronavirus* S1/S2 and S2’ cleavage sites and phylogenetic tree based on S protein sequence. The 50 amino acids regions around the S1/S2 (**A**) and S2’ (**B**) S protein sites were aligned with MAFFT using Geneious 10 software package. An asterisk (*) indicates a position which has a single, fully conserved residue. (**C**) Protein sequences of the complete S protein were aligned using MAFFT within Geneious 10 software package and a maximum likelihood phylogenetic tree of the complete S protein was generated with PhyML. The tree was rooted using MHV-A59. Numbers at nodes indicate the bootstrap support on 100 replicates. Scale bar indicates the estimated number of substitutions per site. Accession numbers for complete spike (S) protein sequences used: CCoV-23/03(AAP72150.1), CCoV-1-71 (AAV65515.1), CCoV-A76 (AEQ61968.1), TGEV-Purdue (ABG89335.1), FCoV-TN406 (BAC05493.1), FCoV-RM (ACT10854.1), FCoV-WSU-79-1146 (YP_004070194.1), FCoV-WSU-79-1683(AFH58021.1), FRCoV-NL-2010(AKG92640.1), MinkCoV-WD1127(ADI80513.1), MinkCoV-WD1133 (ADI80523.1), PEDV-CV777 (AAK38656.1), BatCoV-HKU10 (AFU92104.1), HCoV-229E (BAL45637.1), HCoV-NL63 (AAS58177.1), MHV-A59 (AAA46455.1).

Alignment of protein sequences of various representative alphacoronavirus S1/S2 S cleavage sites reveals a clear demarcation between serotype I and II sequences (fig. 2A). Serotype I CCoV and FCoV all contain a 16-19 amino acid insert, with a stretch of basic arginine (R) and lysine (K) residues (R/K-R-X-R-R), flanked by fairly well conserved N‐ and C-terminal regions. This insert is absent in all serotype II FCoV, CCoV, and TGEV sequences. It is interesting to note that neither ferret, mink, or other alphacoronaviruses contain such insert. This analysis highlights the distinctive feature that appears to be uniquely found in serotype I *Alphacoronavirus 1* strains. The lack of an S1/S2 furin site in CCoV-A76 S protein sequence and its good alignment with serotype II sequences is in agreement with the fact that only the NTD of the CCoV-A76 S protein is serotype I-like with the rest of the protein being serotype II-like, including the region around the S1/S2 site.

Alignment of S protein sequences in the region which includes the S2’ cleavage site and the fusion peptide reveals a very similar partitioning between serotype I and II FCoV and CCoV strains (fig. 2B). In this alignment, the S2’ site demarcates a striking change in amino acid conservation, with only one identical site found upstream of the S2’ cleavage site whereas the region immediately downstream of the cleavage site had 16 identical sites and corresponds to the coronavirus fusion peptide. This analysis also shows the presence of insertions/deletions in the region upstream of the S2’ site. Interestingly, a conserved sequence pattern is observed in serotype II FCoV and CCoV sequences along with TGEV, with the presence of a stretch of basic residues (arginine/lysine) interrupted by a single bulky hydrophobic residue (phenylalanine/tyrosine) upstream of the S2’ site. A less conserved site is found for serotype I FCoV and CCoV, however all are less basic in nature and lack the K-R-K motif observed in all serotype II sequences. In a similar fashion to the alignment at the S1/S2 site, CCoV-A76 S2’ site aligned best with serotype II sequences.

Finally, we performed a phylogenetic analysis based on full-length S protein alignment (fig. 2C). This analysis reveals a different partitioning of *Alphacoronavirus 1* strains compared to our previous analyses. Serotype I FCoV and CCoV clustered in one group and serotype II FCoV, CCoV, and TGEV grouped in another. Furthermore, the CCoV-A76 strain is found at an intermediate position between the two serotype groupings (20), a result that reflects the recombinant nature of its S protein. This phylogenetic analysis clearly demarcates strains according to the previously characterized serotypes and not according to which host species the strains infect, as was observed when performing the analysis with complete genome or ORF1ab sequences. Using such S-based phylogenetic analyses, we propose that the *Alphacoronavirus 1* species be sub-classified as clade A, corresponding to serotype I FCoV and CCoV, and clade B, corresponding to serotype II FCoV and CCoV and TGEV-like viruses.

## Discussion

Current classification within the *Alphacoronavirus 1* species is not well defined and often fails to recognize the profound differences observed between well-established *Alphacoronavirus 1* serotypes. Adding to the confusion are the different terms used to designate various *Alphacoronavirus 1* strains: FCoV serotypes and types; CCoV serotypes, types, and genotypes; and TGEV which is not classified according to FCoV/CCoV serotypes. We propose a more unified classification, based on the important differences between the S proteins of serotype I and II viruses. Our analysis reveals two well-defined clades, clade A and clade B, corresponding to the serotype groupings. Both clades contain FCoV and CCoV strains, while TGEV only belongs to clade B, in agreement with the finding that TGEV is most related to clade B (serotype II) CCoV. We recommend the inclusion of representatives of both clades when performing phylogenetic analysis of alphacoronaviruses. The proposed classification scheme for the alphacoronaviruses is similar to the one used to characterize lineages and clades of avian influenza viruses. Indeed, instead of performing phylogenetic analyses on the entire genomes, avian influenza classifications are based on the surface proteins genes, e.g. hemagglutinin (HA) (24).

In addition to better matching serological and phylogenetic groupings, S-based phylogenies offer other advantages. Because the S gene is frequently shuffled by recombination events, such classifications allows the grouping of viruses that have a shared S gene. In our analyses, using complete genome phylogenies, strains were clustered according to the host they infected whereas the S-based phylogeny allowed to group FCoV and CCoV strains together in different clades. Our approach allows for a better understanding of the complex phylogenetic relationships and evolutionary history observed in coronaviruses. Furthermore, phylogenetic analysis on S proteins can reveal relationships that are not observed using ORF1ab or complete genome-based analysis. In particular, in a study characterizing novel deltacoronaviruses, Woo and colleagues showed that the S proteins of alphacoronaviruses are more closely related to deltacoronaviruses than to other coronavirus genera (3). During our phylogenetic analyses, we took note of a distinct *Alphacoronavirus* S protein sequence from the Asian leopard cat coronavirus (accession no. ABQ39958.1) and noticed its much smaller size (1035 amino acids) compared to other alphacoronaviruses (1466 and 1356 amino acids for FCoV-RM and HCoV-NL63 respectively). Phylogenetically, we observed that it was highly divergent and did not cluster with the other alphacoronaviruses analyzed (data not shown). As only an incomplete genomic sequence was available, it was not included in our analyses.

Our alignment analysis of the cleavage sites of alphacoronaviruses revealed relatively well-conserved functional regions within members of the same clade, but with clear divergence when comparing sequences from two different clades. For example, the presence of a furin motif consisting of the basic stretch of residues R/K-R-X-R-R found only in the S1/S2 site of clade A viruses along with the K-R-K motif found only in the S2’ site of clade B viruses, could be used for rapid determination of clade inclusion in samples which are difficult to sequence at a whole genome level.

## Acknowledgments

We thank members of the Whittaker lab for helpful discussions. This work was funded by a research grant from the Morris Animal Foundation (D10FE-511). Work in the authors’ lab is also funded by the Winn Feline Health Foundation (W15-026) and the Cornell Feline Health Center.

